# Revisiting *Orthotospovirus* Phylogeny Using Whole Genomic Data and a Hypothesis for their Geographic Origin and Diversification

**DOI:** 10.1101/2020.04.07.031120

**Authors:** Anamarija Butković, Rubén González, Santiago F. Elena

**Affiliations:** Instituto de Biología Integrativa de Sistemas (I^2^SysBio), Consejo Superior de Investigaciones Científicas-Universitat de València, 46182 Paterna, Spain; The Santa Fe Institute, Santa Fe, NM 87501, USA

**Keywords:** *Bunyavirales*, phylogenomics, phylogeography, recombination, segmented viruses, segments reassortment, selection, *Tospoviridae*, virus taxonomy

## Abstract

The family *Tospoviridae*, a member of the *Bunyavirales* order, is constituted of tri-segmented negative-sense single-stranded RNA viruses that infect plants and are also able of replicating in their insect vectors in a persistent manner. The family is composed of a single genus, the *Orthotospovirus*, whose type species is *Tomato spotted wilt virus* (TSWV). Previous studies assessing the phylogenetic relationships within this genus were based upon partial genomic sequences, thus resulting in unresolved clades and a poor assessment of the roles of recombination and genome shuffling during mixed infections. Complete genomic data for most *Orthotospovirus* species are now available at NCBI genome database. In this study we have used 62 complete genomes from 20 species. Our study confirms the existence of four phylogroups (A to D), grouped in two major clades (A-B and C-D), within the genus. We have estimated the split between the two major clades ∼3,100 years ago shortly followed by the split between the A and B phylogroups ∼2,860 years ago. The split between the C and D phylogroups happened more recently, ∼1,465 years ago. Segment reassortment has been shown to be important in the generation of novel viruses. Likewise, within-segment recombination events have been involved in the origin of new viral species. Finally, phylogeographic analyses of representative viruses suggests the Australasian ecozone as the possible origin of the genus, followed by complex patterns of migration, with rapid global spread and numerous reintroduction events.

**IMPORTANCE:** Members of the *Orthotospovirus* genus infect a large number of plant families, including food crops and ornamentals, resulting in multimillionaire economical losses. Despite this importance, phylogenetic relationships within the genus were established years ago based in partial genomic sequences. A peculiarity of orthotospoviruses is their tri-segmented negative sense genomes, which makes segment reassortment and within-segment recombination, two forms of viral sex, potential evolutionary forces. Using full genomes from all described orthotospovirus species, we revisited their phylogeny and confirmed the existence of four major phylogroups with uneven geographic distribution. We have also shown a pervasive role of sex in the origin of new viral species. Finally, using Bayesian phylogeographic methods, we assessed the possible geographic origin and historical dispersal of representative viruses from the different phylogroups.

---

The order *Bunyavirales* is full of emerging pathogens that cause severe diseases in humans (*e*.*g*., the Crimean-Congo and hantavirus hemorrhagic fevers, the La Crosse encephalitis or the Rift Valley fever), farm animals (*e*.*g*., Nairobi sheep disease, ovine and bovine Schmallemberg diseases, or Akabame ruminants’ disease) and agronomically important crops (*e*.*g*., tomato spotted wilt, melon yellow spot or peanut bud necrosis diseases). *Bunyavirales* are all arboviruses, and are transmitted by hemato- or phytophagous arthropods. After the last International Committee on Taxonomy of Viruses (ICTV) reclassification (Adams et al., 2017), two families of plant-only infecting bunyavirus have been designated: the *Fimoviridae* and the *Tospoviridae. Fimoviridae* contain a single genus, the *Emaravirus*, and nine recognized species. Likewise, all *Tospoviridae* species were classified within the *Orthotospovirus* genus. Orthotospoviruses have spread around the world causing important loses in vegetable, legume and ornamental crops (Gilbertson et al., 2015). Along with crops, they infect a large variety of weed species; usually each virus species having a wide range of plant hosts. As an example, only for *Tomato spotted wilt virus* (TSWV), the type member of the genus, more than 900 species belonging to over 90 monocotyledonous and dicotyledonous plants have been described as susceptible hosts (Pappu et al., 2009).

The genome structure of orthotospoviruses consists of three negative-sense single-stranded RNAs (Baltimore’s class V; Fig. 1) that differ in size: segments L (large; ∼8.9 kb), M (medium; ∼4.8 kb) and S (small; ∼2.9 kb). Only orthotospoviruses and tenuiviruses (a plant-infecting genus of the *Phenuiviridae* family within the *Bunyavirales*) use the ambisense strategy to produce their mRNAs (Hull, 2014). Segment S encodes for the nucleoprotein (N) in negative-sense and for the protein NS_S_ in positive-sense. Protein NS_S_ is involved in the suppression of plant RNA silencing (Hedil and Kormelink, 2016). The segment M encodes for a precursor polyprotein of the glycoproteins G_N_ and G_C_ in negative-sense. In positive-sense it encodes for the protein NS_M_, which is responsible for the modification of plasmodesmata that enable the infectious nucleocapsid units to move throughout, allowing intercellular movement (Storms et al., 1995). Singh and Savithri (2015) showed that NS_M_ also remodels the endoplasmic reticulum networks to form vesicles to facilitate viral replication and movement. The segment L encodes for the RNA-dependent RNA polymerase (RdRp) in negative-sense.

**FIG 1.**
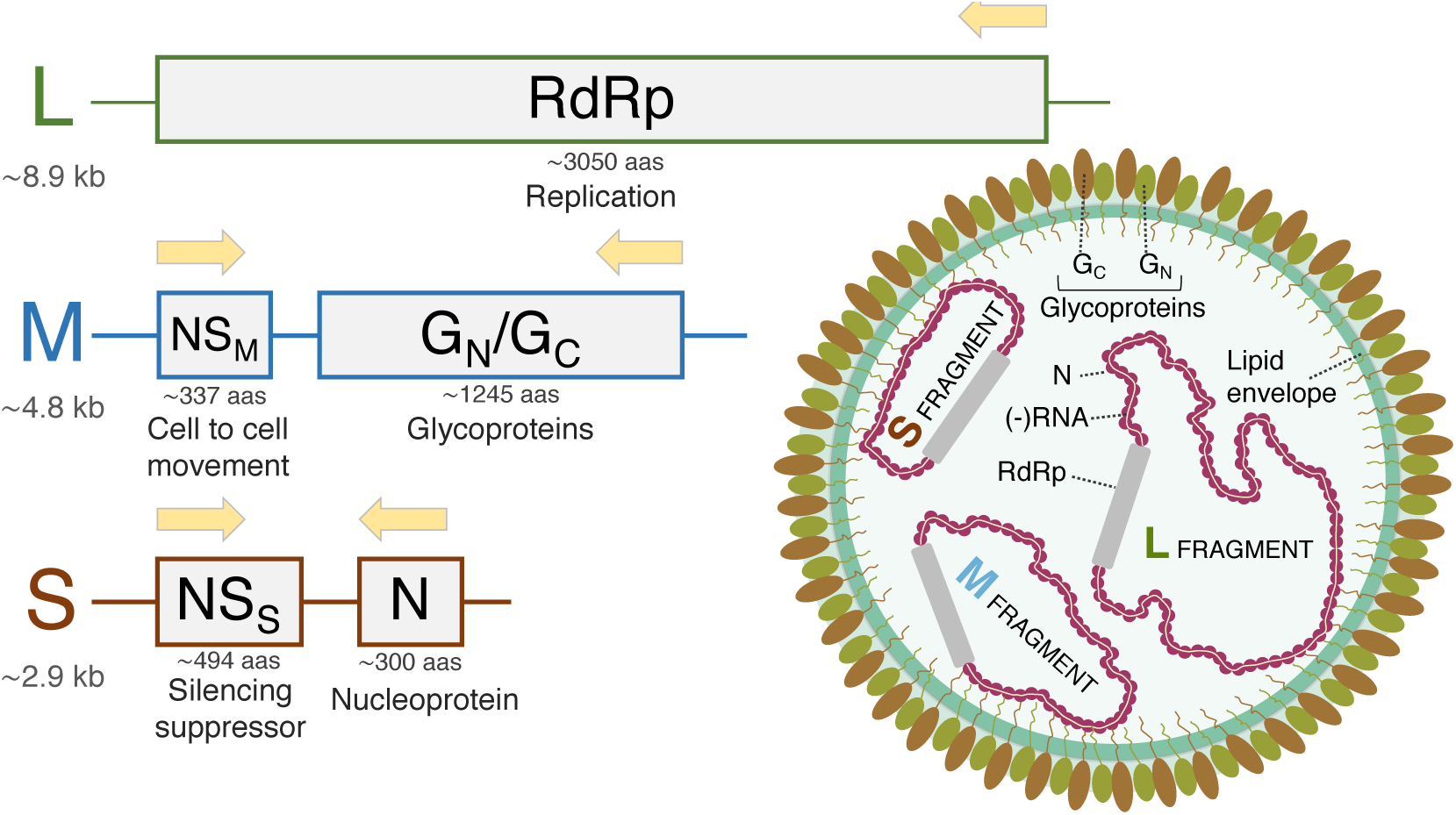
Schematic representation of the *Orthotospovirus* tri-segmented ambisense genome. The main function, and length of each protein is indicated. The translation sense of each ORF is indicated by arrows.

*Orthotospovirus* particles are spherical with a diameter of 80 - 110 nm. The particles are made of a lipid envelope that is spiked by two glycoproteins, G_N_ and G_C_ (Fig. 1). These glycoproteins form heterodimers that compose oligomeric structures, the exact conformation they make on the outside of the envelope has not been determined experimentally yet. The cytoplasmic tails of the glycoproteins interact with the nucleoproteins that are inside the envelope (Ribeiro et al., 2009). The nucleoproteins encapsidate the three viral RNA segments forming the ribonucleoprotein complexes. These complexes are associated with the RdRp. The number of ribonucleoprotein complexes packed into each virion remains unknown, but for being viable, virions must at least include one copy of the three genomic segments.

*Orthotospovirus* particles are transmitted between susceptible plants through thrip vectors (Rotenberg et al., 2015). Up to fifteen species of common thrips have been reported as facultative vectors (Rotenberg et al., 2015). All these vectors belong to the *Frankliniella* and *Thrips* genera (*Terebrantia* suborder, *Thysanoptera* order, *Thripinae* subfamily, *Thripidae* family). The transmission of orthotospoviruses by thrips has several unusual characteristics. Only the first and second larval instances can acquire the virus and this competence decreases with age. Although thrips can retain lifelong infectivity, their ability to transmit the virus can be erratic (Hull, 2009). The ability to retain the infectivity is due to the mode of transmission which is propagative, meaning that the virus is able to replicate in its vector (Rotenberg et al., 2015; Fletcher et al., 2016). As the virus replicates in the thrips, with some of the thrips (*e*.*g*., *Frankliniella occidentalis*) being extreme polyphagous, there are chances that mixed infections of orthotospoviruses occur both within the insect and/or the plant. That would lead to recombination, segment reassortment, and component capture events. The global spread of thrips and orthotospoviruses could lead to global disease outbreaks (Gilbertson et al., 2015). Recent studies have shown that plant hosts infected with orthotospoviruses can affect the feeding behavior and even the survival of the vector (Sundaraj et al., 2014). Specifically, Shalileh et al. (2016) showed how TSWV manipulated *F. occidentallis* via the host plant nutrients to enhance virus transmission and spread.

In this work, we have reevaluated the phylogenetic status within the *Orthotospovirus* genus based on available full genomes. Furthermore, to identify events of heterologous genome reassortment during mixed infections as well within-segment recombination events, phylogenies were done for each one of the three segments and for each one of the five proteins and incongruences among trees were evaluated. All the phylogenies were done using Bayesian methods. Along with the phylogenies there were other evolutionary characteristics of the viruses that were analyzed: selection pressures, recombination events and phylogeographic distribution and dynamics. Based on these results, we have drawn a plausible scenario for the geographic origin and ulterior spread and diversification of *Orthotospovirus*.

## RESULTS AND DISCUSSION

### Pervasive evidences of recombination and segment reassortment analyses

Prior to any other study, we evaluated the role of recombination during the phylogenetic diversification of the orthotospoviruses (File S1). The PHI test for the protein alignment found evidences of recombination between the *G*_*N*_*G*_*C*_ (*P* = 0.0053) and *NS*_*M*_ (*P* = 0.0017) genes in segment M; consistently giving also significant values for segment M as a whole (*P* < 0.0001). One possible mechanism could be the physiological role of NS_M_, responsible for viral movement and regulation of apoptosis. In different plant hosts there are different proteins involved in binding to NS_M_ (Knipe and Howley, 2013). Differences in NS_M_ structure could have a major impact in cell-to-cell transportation. It has also been suggested that the NS_M_ protein is an evolutionary adaptation of an insect-only infecting bunyavirus to a plant host, where it determines symptomatology and host range (Tsompana et al., 2004). Therefore, when new viruses spill into new hosts, often in mixed infections with other well-adapted viruses, within segments recombination or segments reassortment events may give rise to new viruses with enhanced movement in the novel host.

Furthermore, *G*_*N*_*G*_*C*_ encodes for the precursor of the two glycoproteins G_N_ and G_C_, which is consistent with the GARD algorithm detecting a recombination breakpoint between them. This suggest independent evolutionary histories for the two glycoproteins. It is known that G_N_ is responsible for attachment to epithelial cells in the midgut of thrips as a mode of more effective transmission (Knipe and Howley, 2013). Recombination between these two proteins can readily generate new viruses that would explore their potential of transmission by novel thrips species. The exact breakpoint between *G*_*N*_*G*_*C*_ was estimated using GARD to be between amino acid positions 768 and 769. This partition will be later used in all the BEAST analyses described in the following sections. Further supporting the recombinogenic nature of orthotospoviruses, the pairwise comparison of the MCC trees obtained for the three segments (Fig. S1) using tanglegram in Dendroscope was incongruent between all segments. Mainly, the incongruence in tree topologies was entirely driven by TSWV.

In conclusion, recombination within or reassortment of M segment during mixed infections may create new viruses with improved movement in more hosts or transmission ability using new vectors.

### Full genome phylogenetic reconstructions confirm the existence of four phylogroups

Maximum clade credibility (MCC) Bayesian trees were generated from 62 full genomic sequences belonging to 20 different orthotospovirus species (Fig. 2). The trees were built with the heterogeneous model configuration described in Material and Methods section that introduces a partition between segments and, in the case of segment M, it accounts for the existence of a recombination breaking point within the glycoprotein gene. The MCC obtained from the full-genome shows a significant clustering into four different phylogroups A, B, C, and D. Pappu et al. (2009) defined phylogroups in terms of geographical origin of the viruses. However, our full-genome analysis of a larger and updated dataset does not support the notion of independent geographic origins for each phylogroup. Instead, our phylogroups contain viruses from diverse origins. For instance, Pappu et al. (2009) so-called American phylogroup is part of what we have defined here as phylogroup A that contains a mixture of viruses isolated from Asia, Europe, Australia, and America. Likewise, Pappu et al. (2009) Eurasian phylogroup, contains viruses from what we have defined here as phylogroups C and D. Phylogroup D includes an isolate of *Capsicum chlorosis virus* (CaCV) from Australia. We only incorporated a full-genome sequence from Africa since other sequences from this region failed to meet the criteria mentioned in the Materials and Methods section. It is worth mentioning that most of orthotospoviruses have a broad worldwide distribution, though many genomes have not been fully sequenced and assembled after identification, thus limiting the number of full-genomes available for our study. We expect that future accumulation of additional full-genome sequences for some species would modify the number of clades in the phylogeny shown in Fig. 2. In fact, it has been suggested using partial sequences that *Groundnut chlorotic fan-spot virus* (GCFSV) and *Groundnut yellow spot virus* (GYSV) would likely establish a different clade (Oliver and Whitfield, 2016).

**FIG 2.**
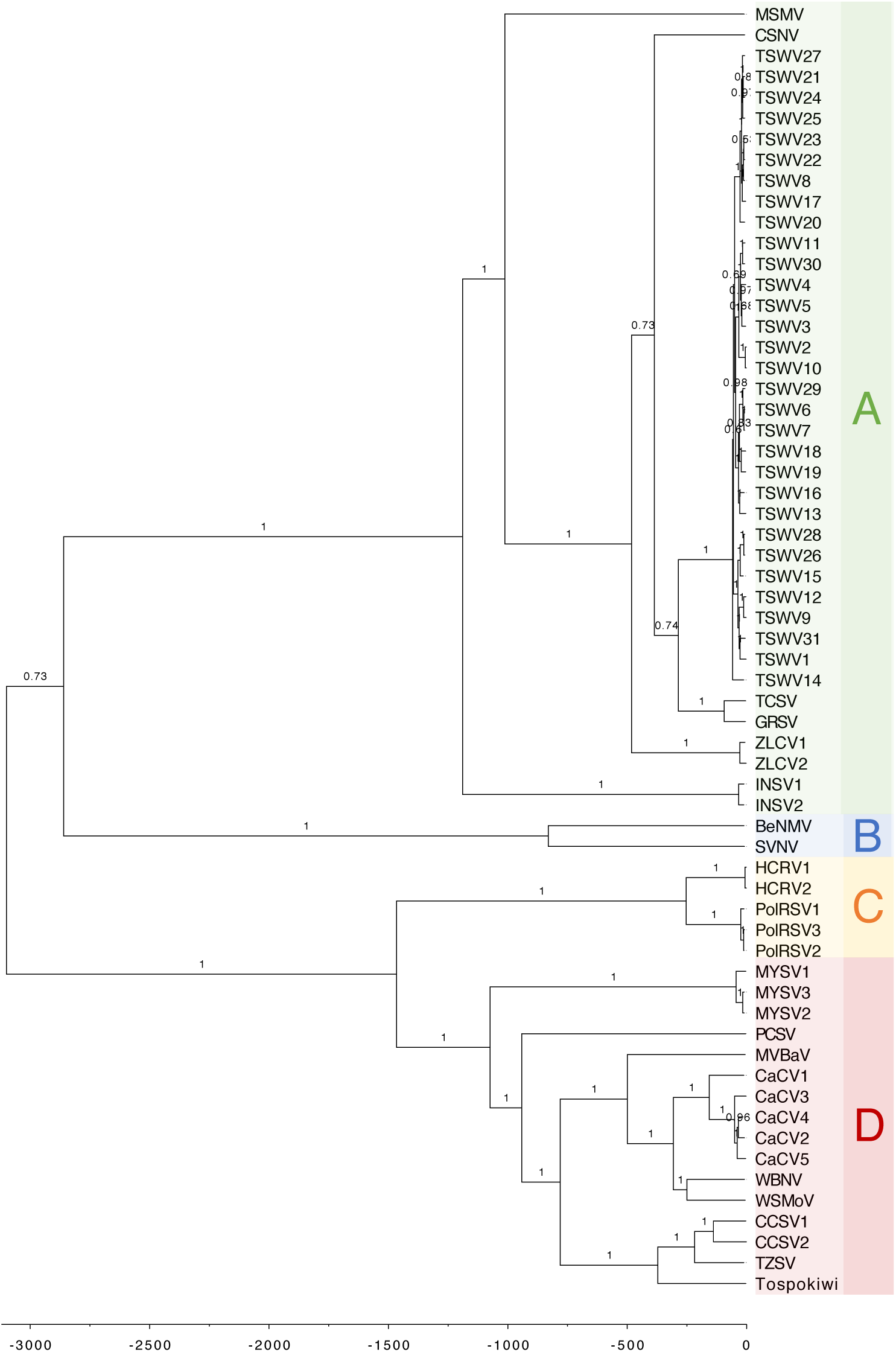
MCC tree obtained from the concatenated full genome. Phylogroups A, B, C, and D are indicated with different color boxes. Branch labels indicate *BPPs*.

In general, most nodes in the MCC tree had a high statistical support (0.95 < *BPP* ≤ 1). However, a few nodes had lower support. The node with the lowest support (*BPP* = 0.52) corresponded to the divergence of TSWV strains 22 and 23; while other nodes within the TSWV clade also had *BPP* < 0.70 support. Two other clades had intermediate support values (*BPP* ∼ 0.70): the basal node in which phylogroups A and B split apart, and the node leading to the speciation events between *Groundnut ringspot virus* (GRSV), *Tomato chlorotic spot* (TCSV), TSWV, and *Zucchini lethal chlorosis virus* (ZLCV).

### Rates of molecular evolution are highly heterogeneous among genes and lineages

Estimated rates of molecular evolution ranged between 10^−4^ - 2×10^−3^ substitutions per site and per year for segment L, between 6×10^−4^ - 10^−3^ for segment M and between 9×10^−4^ - 1.7×10^−3^ for segment S (color legends in Fig. S2).

Heterogeneity in rates of molecular evolution among viruses (and strains of the same virus) have been observed for the different genes (Fig. 3). For the *RdRp* gene, the slowest estimated rates of evolution corresponded to all the nodes, while the fastest corresponded to branches separating the phylogroups A-B and C-D. It showed the slowest rate of evolution amongst the five genes.

**FIG 3.**
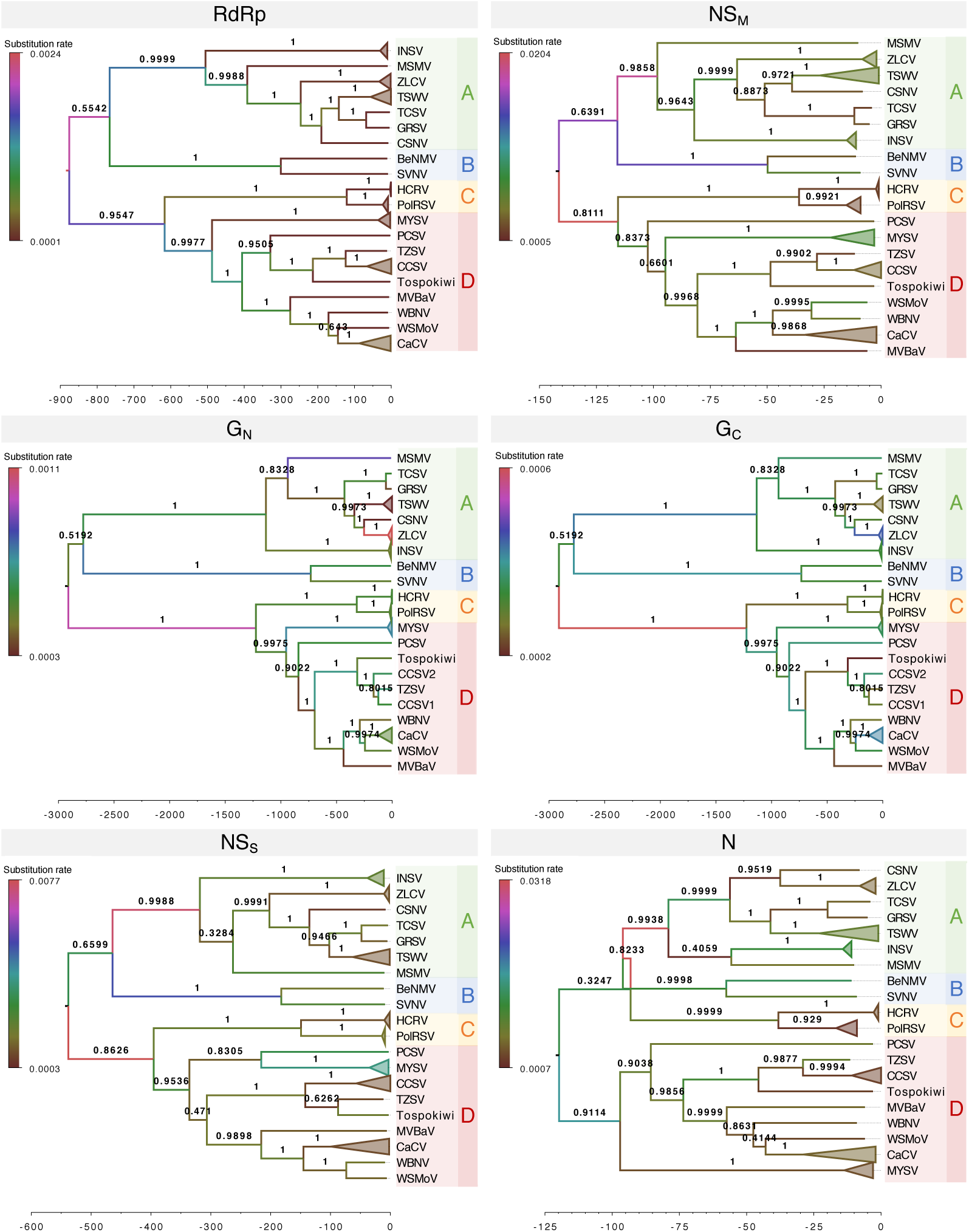
Six MCC trees of the 20 *Orthotospovirus* species used in this study. The trees were obtained from each individual gene protein sequence. Phylogroups A, B, C, and D are indicated with different color boxes. Numbers above branches represent *BPPs*. Viral species with more than one isolate has been collapsed (triangles). Branches of different color indicate differences in rates of molecular evolution according to the left scale. The scale below the trees represents years.

For gene *NS*_*M*_ the fastest rates of molecular evolution corresponded to the basal branches leading to phylogroups A-B and C-D, with branches leading to phylogroups A and B. By contrast, the slowest evolutionary rates for *NS*_*M*_ were estimated for the basal branch leading to the phylogroup C, and to terminal branches leading to CaCV and *Mulberry vein banding virus* (MVBaV).

Regarding the glycoprotein genes, for gene *G*_*N*_, we observed fast rates of molecular evolution in the basal branches leading to clades A-B, C-D and phylogroups A, B, C and D. The fastest evolutionary rates were for ZLCV, *Melon severe mosaic virus* (MSMV) and *Melon yellow spot virus* (MYSV). The slowest evolving branches were for TSWV and MVBaV. For gene *G*_*C*_, the fastest rate of molecular evolution corresponds the branch leading to the phylogroups C-D. By contrast, the slowest rates of molecular evolution correspond to the basal branch leading to the yet to be named tospovirus isolated from kiwi plants (hereafter referred as Tospokiwi) and TSWV.

Overall, the *NS*_*S*_ gene showed slow rates of evolution in most basal branches, except those leading to the phylogroup A and B and also phylogroups C-D. MYSV showed faster evolutionary rate compared to the other viruses.

High rates of molecular evolution for the *N* gene were the basal branches leading to phylogroups A and D. The slowest rates observed for the *N* gene were in the majority of basal branches except the tip branches leading to *Bean necrotic mosaic virus* (BeNMV) and *Impatiens necrotic spot virus* (INSV).

These results suggest a pattern of variable rates of evolution along the diversification of orthotospoviruses. Overall, accelerated rates of evolution have been observed in all genes in the origin of phylogroups A-B and C-D, in some internal branches within phylogroups leading to speciation events and in some terminal branches giving rise to novel strains of a particular virus. Even for fast evolving lineages, differences among genes exist. For example, MYSV showed very fast rates of molecular evolution in the *G*_*N*_ and *NS*_*S*_ genes but a rather slow one in the other genes.

### Estimation of divergence times

As it can be shown in Fig. 2 for the full genome, the time of the diversification between clades A-B and C-D was estimated to be 3099 years ago (ya) (95% HPD confidence interval: 5192 - 1758 ya) (*BPP* = 1). Soon after, ∼2860 ya (95% HPD confidence interval: 4772 - 1605 ya), the A-B clade diverged into phylogroups A and B (*BPP* = 1). The C-D clade diverged into phylogroups C and D (*BPP* = 1) around 1465 ya (95% HPD: 2421 - 854 ya).

In addition, MCC trees were also generated for the individual segments (Fig. S2) and individual genes (Fig. 3), with estimates of divergence times generated from these partial trees being consistent with those obtained for the full genomes. For example, in the case of the three segments (Fig. S2), the estimated divergence of phylogroups A-B and C-D from each other was in the range of 2786 - 875 ya, whereas the divergence between clades A-B and C-D took place between 2621 - 616 ya. Genes *N* and *NS*_*M*_ have more recent divergence times, estimated about 121 (95% HPD: 548 - 36 ya) and 141 (95% HPD: 366 - 45 ya) between phylogroups A-B and C-D, respectively. Divergence times between the four mayor phylogroups estimated for gene *NS*_*S*_ is 538 ya (95% HPD: 2519 - 81 ya). However, estimates obtained using the *G* gene place the divergence between A-B and C-D clades much earlier on time, 2911 ya (95% HPD: 7010 - 1122 ya).

### A phylogeographic analysis of the origin and dispersal of *Orthotospovirus*

For these analyses, instead of using countries, or even continents, as geographic units, we opted to use more biologically relevant units: the eight biogeographic realms, or ecozones, into which World Wide Fund for Nature (WWF) divides the world (Schulz, 2005). These ecozones consider the geological barriers that kept plants and animals separated from other areas, having a shared evolutionary history. Seventy-three sequences from a reduced dataset of all N proteins available have been used for these analyses. The results are shown in Fig. 4. Australasia (Australia, New Guinea and neighboring islands) has been identified as the most likely origin of orthotospoviruses (*RSPP* = 0.4961), with the second highest supported origin being the Western Palearctic (Western Europe and North Africa) (*RSPP* = 0.3929). However, this conclusion has to be taken with some caution, since Pappu et al. (2009) and Oliver and Whitfield (2016) had suggested that TSWV isolates were actually imported into Australia from Europe ca. 1915. The second highest support was for Western Palearctic. Eurasia, as part of the Palearctic ecozone, contains the largest number of different *Orthotospovirus* species, giving more plausibility to the hypothesis of the Palearctic ecozone as the possible origin of the genus.

**FIG 4.**
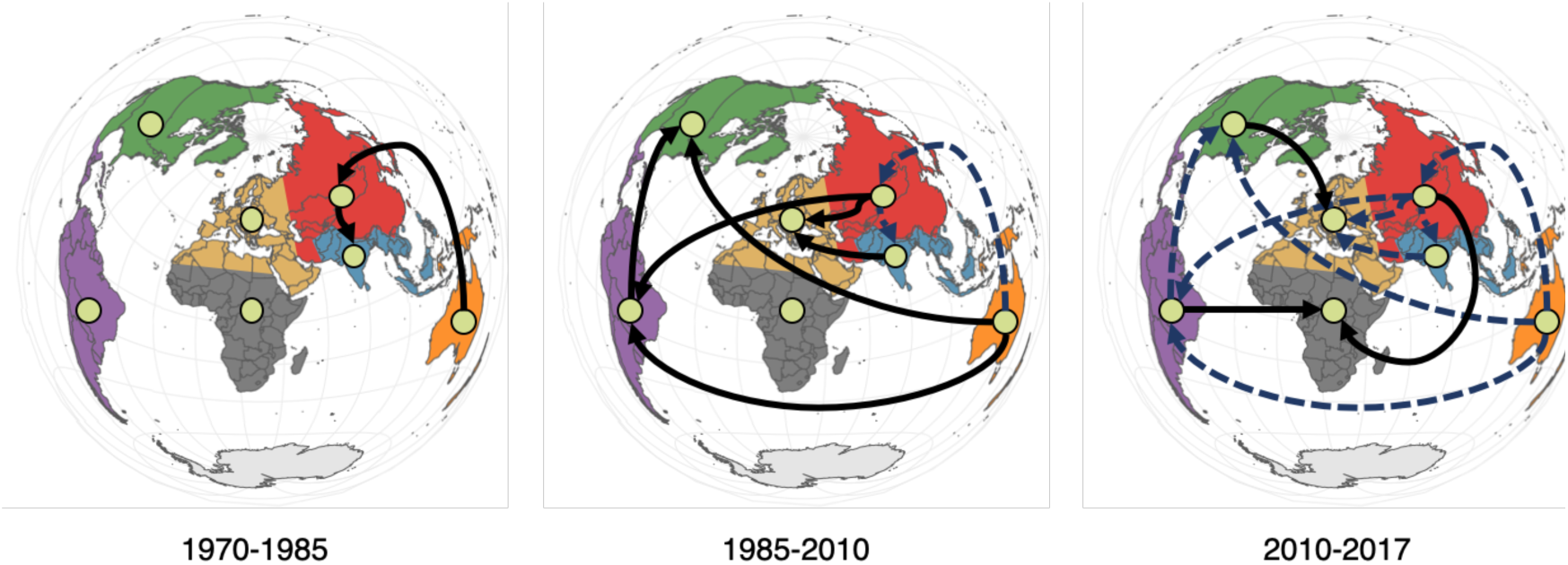
Phylogeographic representation of the migration pattern of the 73 orthotospovirus species based on the analysis of the *N* gene. The colored regions represent different ecozones: Afrotopic in dark grey, Australasia in orange, Indomalayan in blue, Nearctic in green, Neotropical in purple, Eastern Palearctic in red, and Western Paleartic in yellow, respectively. Arrows show the direction of the movement, with the black continuous lines representing movements in the current period of time and blue dashed one’s past movements.

From Australasia, viruses were introduced into the Western Palearctic and from there into Indomalaya (South Asian Subcontinent and Southeast Asia). Afterwards these three ecozones have been stablished as dispersion centers to colonize the rest of the ecozones; Neotropic (South America and the Caribbean), Nearctic (North America), Eastern Palearctic (Eurasia) and Afrotopic (Sub Saharan Africa) in that order, with numerous reintroductions. It is interesting that the sequences that were introduced numerous times in Eastern Palearctic and Afrotropic became later isolated. The BSSVS analysis shows the strongest support for the dispersal route from the Indomalaya ecozone into the Eastern Palearctic one (*BF* = 236.2).

We also tested the null hypothesis of a random distribution of geographic origins on the MCC tree generated with the full genome (Fig. S3) using Treebreaker (Ansari and Didelot, 2016). The original definition of phylogroups was based on the geographic origin. Therefore, the alternative hypothesis would be that geographic origins must be differentially distributed among the four phylogroups. The branches on the MCC tree that successfully reject the hypothesis of random distribution are for phylogroups B, C, D, and for viruses TSWV and INSV. Phylogroup B has sequences from America, phylogroup C has sequences from Asia and Europe while the phylogroup D has sequences mostly from Asia. TSWV is found all around the globe but the sequences we used in this study predominantly came from Asia. In the case of INSV we have sequences from Europe. We can see that all the sequences coming from Asia and Europe show a nonrandom distribution of geographic origin in the MCC tree. Only two viruses coming from America, BeNMV and *Soybean vein necrosis virus* (SVNV) show significant support for a geographic correlation. Interestingly, this association must be entirely due to the limited geographic distribution of the vector species (Rotenberg et al., 2015). BeNMV and SVNV are transmitted by the aphid *Neohydatothrips variabilis* which is restricted to Central and North America. INSV came from Europe and is transmitted by *Frankinella spp*. which is found all around the world. Phylogroup C shows strong correlation for viruses *Polygonum ringspot virus* (PolRSV) and *Hippeastrum chlorotic ringspot virus* (HCRV) which come from the Paleartic ecozone and are transmitted by *Thryps spp*. and *Dictyothrips spp*., mostly distributed across Eurasia. Phylogroup D, mostly contains viruses isolated from Asia and is transmitted by *Thrips spp*. which is found everywhere.

### TSWV geographic origin and pandemic dispersal

TSWV is the most represented virus in our dataset and hence provides the richest information to perform phylogeographic analyses for an individual virus. As in the previous section, we have used the WWF ecozones. We have performed two independent phylogeographic analysis of the N protein sequences from TSWV, one based in the entire 229 isolates dataset and a second one based on a reduced dataset of 87 isolates that minimizes overrepresentation of sequences from certain countries (see Materials and Methods). For both datasets, the most likely origin of TSWV was within the Australasia ecozone (*RSPP* = 0.9426 for the reduced dataset and *RSPP* = 0.9016 for the full dataset). However, this conclusion has to be taken with some caution for the reasons discussed in the previous section. In any case, the two data sets differ in the spread of isolates out of Australasia. The results for the reduced dataset are shown in Fig. 5a. In the case of the reduced dataset TSWV isolates emigrated from Australasia first into the Neotropic and then into the Nearctic and the Western and Eastern Palearctic ecozones as recently as 1994 - 1999. Even more recently, around 2007, a second introduction from the Nearctic into the Eastern Palearctic took place. In 2015 several possible parallel introductions into the Afrotropic ecozones from the Neotropic and Easten Palearctic ones are inferred from the dataset, though the strongest supported route was from the Palearctic (*BF* = 215.4). The sequences from the Western Palearctic and Afrotropic after the introductions became isolated.

**FIG 5.**
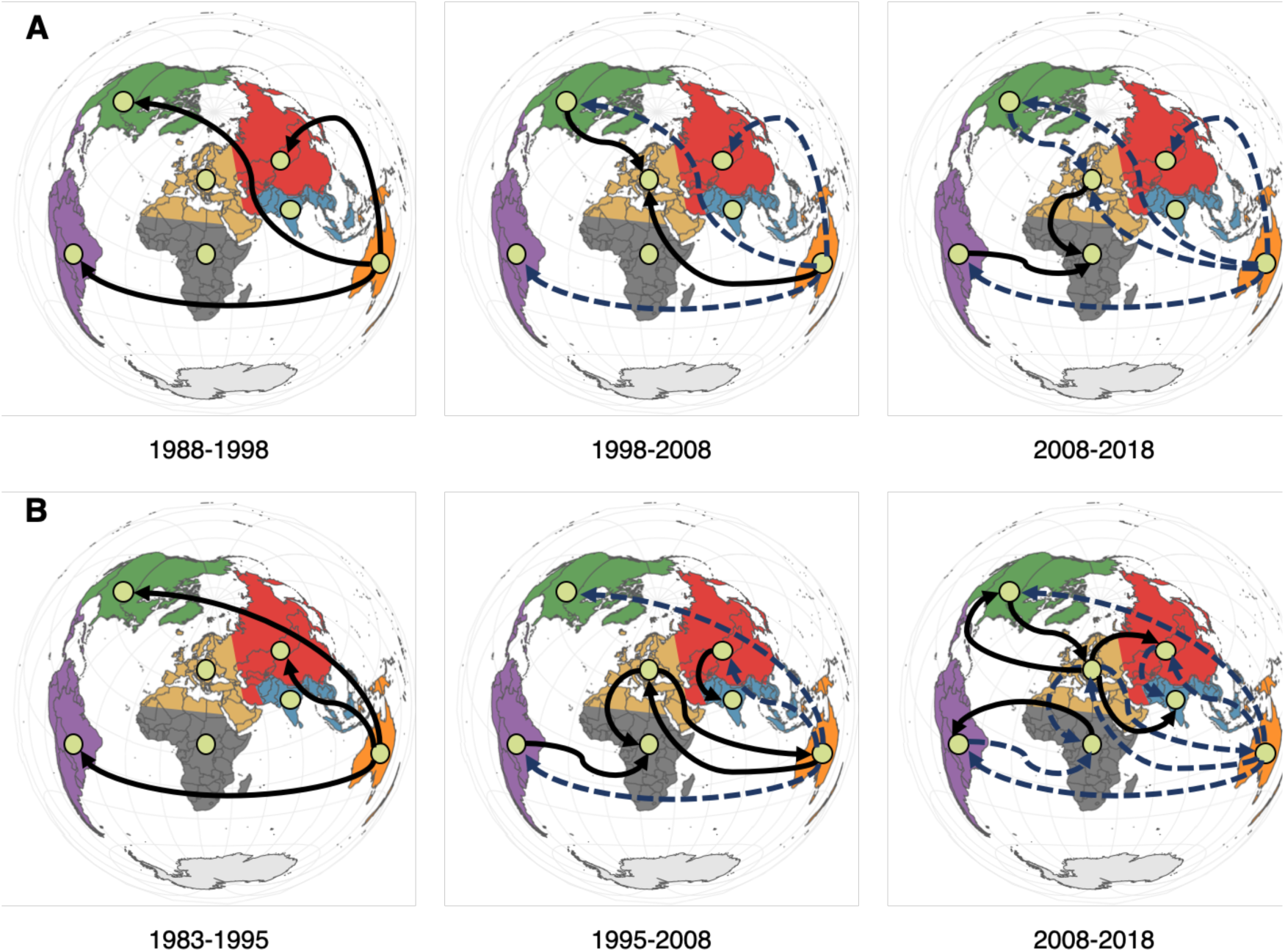
Phylogeographic representation of the migration pattern inferred for TSWV based in the *N* gene. (A) Reduced dataset consistent in 87 sequences. (B) Large dataset composed with 229 sequences. Colors represent different ecozones while lines represent movements, as described in the legend of Fig. 3.

When looking into the phylogeographic diffusion patterns inferred from the full dataset (Fig. 5b), Australasia still appears as the origin of TSWV pandemic as previously mentioned. From there, the virus spread out into the Nearctic, Western Palearctic, Neotropic, and Eastern Palearctic ecozones within less than ten years (between 1988 and 1996). From the Eastern Palearctic, Western Palearctic and Neotropic, isolates were introduced back into the Australasia, Indomalaya (South Asian Subcontinent and Southeast Asia) and Afrotropic ecozones in early 2000s (Fig. 5b). Subsequent reintroductions into Afrotropic were originated from the Eastern Palearctic and Neotropic ecozones (Fig. 5b). In the last decade, isolates from the Afrotropic have diffused into the Neotropic and Indomalaya ecozones (Fig. 5b).

In both sequence datasets the Nearctic, Western Palearctic and Eastern Palearctic ecozones show the largest number of viral lineages. The strongest supported routes in this case were between the Neotropic and Australasia and the Eastern Palearctic and Australasia ecozones (in both cases, *BF* = 226.4). Sequences introduced into the Indomalaya ecozone became isolated after the introduction.

### Variations of population sizes over time

Population sizes over time plots are a useful tool to illustrate overall patterns of genetic diversification throughout time (Rambaut et al., 2018), *i*.*e*. the number of lineages observed on a tree with respect to time. If diversification happens in a constant steady-state manner through time, a straight line is expected when plotting the number of lineages in a logarithmic scale as a function of time. In one hand, if the observed line lays above the null hypothesis straight line, the diversification rate increases with time. In the other hand, if the observed line goes below the straight line, it can be concluded that diversification rate is decreasing with time. In our case, lineages will be considered as different virus species.

In the Skyride plot from the TSWV analysis of the reduced (87 sequences) dataset (Fig. 6a) we can see a rise from 1995 until 2002, where we see a slight decline from 2003 until 2005. From 2005 until 2012.5 we have a slight rise after which we have a steady population size until 2018 (Fig. 6a). While the Skyride plot of the full TSWV sequence set (229 sequences) shows a similar pattern with the exception of a decline in population size between the years 2005 – 2012.5 and a decline after that until 2018 (Fig. 6b). For both datasets, the estimated median effective population size was 600 individuals.

**FIG 6.**
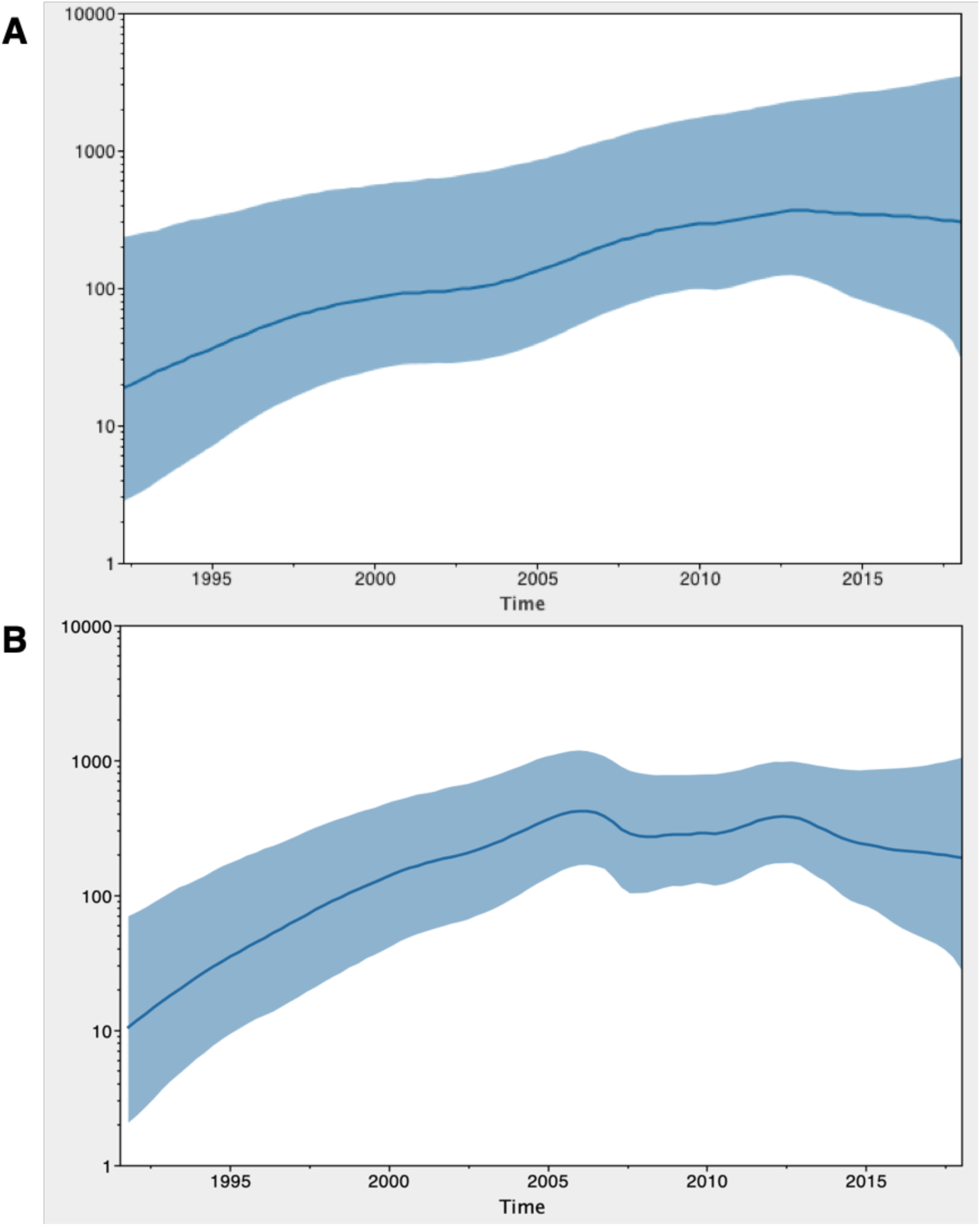
Population sizes over time plots generated for TSWV. (A) Reduced dataset consistent in 87 sequences. (B) Large dataset composed with 229 sequences. The ordinate axis represents the number of lineages (in logarithmic scale), while the abscissa axis represents time from the origin. Light blue lines represent the 95% HPD confidence interval. Tree model root height 95% HPD for the full sequence set: 26 - 32 and for the reduced set: 26 - 39.

### Positive selection analysis

Selection analysis have been performed with the MEME and aBSREL algorithms. Codons under positive selection have been found in all the protein coding sequences except *N* (File S2): position 26 (*P* = 0.05) of NS_M_, positions 468 (*P* = 0.01), 476 (*P* = 0.05), 479 (*P* = 0.05), and 490 (*P* = 0.01) of NS_S_, positions 4 (*P* < 0.01) and 813 (*P* < 0.01) of G_N_G_C_, and positions 1066 (*P* = 0.03) and 1728 (*P* = 0.04) of L (File S2 and Fig. S4) were all under significant positive selection. The algorithm aBSREL indicated nodes under positive selection for all genes except *N* (File S2). These findings point to the fact that beneficial mutations arose in the four proteins and drifted to fixation in the viral population, resulting in adaptation to new hosts and/or new vectors.

This change in the nucleic acid content and the worldwide spread of the thrips vectors, especially *F. occidentalis* (Knipe and Howley, 2013), could lead to the spread of some lineages that had beneficial mutations for adaptation to new hosts-vectors-environment. This adaptive evolution could explain the fast and worldwide spread of orthotospoviruses during the 1960s and 1970s by the vector *F. occidentalis* (Knipe and Howley, 2013). Reflecting adaptation of the orthotospoviruses and ulterior diversification into new plant hosts used by this very polyphagous vector.

The N protein does not show evidences of positive selection; instead, it shows many instances of significant negative selection (File S2). This could be due to very stringent structural constraints given its multifunctional nature. N is involved in transcription and translation of viral RNA. It has been shown that a single point mutant of *Bunyamwera virus* (BUNV) was either defective for transcription but not translation and *vice versa* (Eifan and Elliot, 2009; Walter et al., 2011). This observation suggests that N may have two functional domains; one involved in transcription and other in translation, each interacting with different cellular cofactors. Since the small N protein forms the ribonucleocapsid complex with viral RNAs and tightly interacts with the RdRp to initiate RNA synthesis, protecting the nascent RNAs from degradation (Knipe and Howley, 2013), even a small structural change might disrupt its function, jeopardizing completion of the viral replication cycle. But considering essential roles of other proteins that show signatures of positive selection this situation in N could also be due to the lack of statistical power of the methods used.

## CONCLUSIONS

The taxonomic status of the plant-infecting bunyaviruses was recently revisited by ICTV. They changed the status of tospoviruses from the genus level to the family level, *Tospoviridae*. The new family only contains, so far, a genus, the *Orthotospovirus*. However, Oliver and Whitfield (2016) have suggested that the family could be classified into five distinct phylogenetic clades whose type members would be GYSV, IYSV, SVNV, TSWV, and WSMoV. This classification is fully consistent with our finding of four phylogroups; TSWV clade would be the representative species of phylogroup A, SVNV of phylogroup B, IYSV would represent phylogroup C, and WSMoV phylogroup D. No full genome sequences for the two viruses from the GYSV clade were available at NCBI, and hence are missing in our analyses. Additional full genome data for the two GYSV clade members would be necessary to confirm if they form an additional E phylogroup as well as to explore their phylogeographic history in relation with the other phylogroups.

Based in partial genome sequences and a smaller number of species, Pappu et al. (2009) classified orthotospoviruses according to their geographic origin in two geogroups: Asian and American. Our analyses suggest a more complex picture and that classification of orthotospovirus species cannot be done based on geographic criteria. Comparing this geographic classification with our genome-based phylogroups, we observe that phylogroups A and B would be mainly of Asian origin while phylogroups C and D would be mainly of American origin, though the geographic distinction is diffuse. Genome reassortment, recombination and selection all played a role in the origin and diversification of orthotospoviruses.

Looking at the results of the phylogeographic distribution of TSWV nucleocapsid sequences and 73 orthotospovirus nucleocapsid sequences we suggest that the origin of orthotospoviruses is the Australasia ecozone, afterwards radiating into other ecozones around the globe. The geographic distribution of vectors correlates well with the geographic distribution of different Orthotospovirus species. Phylogroups B, C, D and TSWV and INSV are transmitted by aphids that are specific for the Neartic/Neotropic and Paleartic regions and are the predominant vectors for each group.

This work is the first phylogenetic analysis done on the family *Tospoviridae* that used well annotated full genome sequences. In line with the results from other studies, we propose a hypothesis that the genus *Orthotospovirus* could reasonably be split into at least four new genera.

## MATERIAL AND METHODS

### Viral species, sequence extraction and database curation

All available full *Orthotospovirus* genomes were obtained from the NCBI Genome Database (www.ncbi.nlm.nih.gov/genome; Brister et al., 2015) in May 10^th^, 2018. The genome collection was further expanded for all orthotospoviruses that were defined as a species with the same criterion used to search the NCBI Genome Database: the complete genomes had to come from the same research paper and had to be sequenced from the same extraction sample. Our working dataset is thus composed of the amino acid sequences derived from 62 full viral genomes (the entire proteome) with well-defined segments and genes belonging to 20 different *Orthotospovirus* species (File S1). All nucleotide sequences were extracted by their accession number with the help of *E*-utilities from Entrez Direct. It was confirmed by further inspection with Geneious Prime version 2019.0.3 (www.geneious.com) that each of the 62 genomes corresponded, indeed, to a different *Orthotospovirus*.

To examine the phylogeographic origins of TSWV and all orthotospoviruses we collected 229 sequences for TSWV (including partial sequences) and 73 for all orthotospoviruses after filtering for duplicates. In the case of all orthotospovirus N proteins we did not use the expanded set because it did not converge which could be caused by problems with the priors or a low temporal signal (*R*^2^ = 0.13). We also created a subset of TSWV sequences where we tried to minimize the bias of overrepresentation from certain countries and from specific dates, we also removed partial sequences. The subset was guided by CD-HIT (Fu et al., 2012) where we clustered the sequences coming from the same ecozone with sequence identity cut-off of 95%, taking into consideration only the ecozones that had the largest number of sequences. After removing the bias and partial sequences in TSWV set, 87 sequences were retained.

### Alignment and assembly of segments and full genomes

All alignments used in this study were generated using MAFFT version 7 (Kazutaka and Standley, 2013) and visually inspected. We generated the following alignments. Firstly, the protein sequences of the five viral genes (*G*_*N*_*G*_*C*_, *N, NS*_*M*_, *NS*_*S*_, and *RdRp*; Fig. 1) were aligned individually. Secondly, to build the alignment of segments, we respected the order of genes in NCBI records and combined *NS*_*M*_ and *G*_*N*_*G*_*C*_ genes into segment M and *NS*_*S*_ and *N* genes into segment S (Fig. 1). Thirdly, we generated the concatenated full viral genomes by combining the three segments in the arbitrary L-M-S order. Segment and full genome joining were performed with FaBox alignment joiner available online (Villesen, 2007). The resulting globally aligned proteome was 5,401 amino acids long, had 966 identical sites and a pairwise identity of 62.3%.

### Tests of phylogenetic congruence and detection of segment reassortment and recombination events

To seek for heterologous segregation of segments in the origin of novel orthotospoviruses, segment trees were compared to one another (M-L, M-S and S-L) using the tanglegram function in Dendroscope version 3 (Huson and Scornavacca, 2012). Recombination analysis within genes and segments was performed using the PHI test implemented in SplitsTree4 (Huson and Bryant, 2006) for recombination on the protein and codon sequences encoded by each gene and concatenated segment. The genetic algorithm for recombination detection GARD (Kosakovsky Pond et al., 2006), as implemented in the www.datamonkey.org server (Weaver et al., 2018) was also used to detect recombination breakpoints in the codon aligned nucleotide sequences of the five *Orthotospovirus* genes.

### Selection of the best models of amino acid substitutions

ProtTest 3 (Darriba et al., 2011) was used to assign the best fitting models of protein evolution for the individually aligned protein sequences based upon the Bayesian Information Criterion (*BIC*) minimal score. For the aligned TSWV N sequences used for the phylogeographic analyses, we used the second best-scoring model, FLU+G, since the best-scoring one, HIVb (Korber, 2000), was not implemented in BEAST version 1.10.4 (Suchard et al., 2018).

### Phylogenetic and phylogeographic trees reconstruction

The maximum likelihood trees of the protein alignments for genes, segments and full genome sequences were generated with IQ-Tree (Nguyen et al., 2013). In the maximum likelihood tree reconstruction, we used the model assigned with ProtTest 3 and 1000 bootstrap replicates. These trees were later used as input for the program TempEst (Rambaut et al., 2016) where we examined the temporal signal and clock-likeness of our data before proceeding to the Bayesian analysis. All the trees for genes, segments, full genome and the phylogeographic analysis had a positive correlation coefficient and significant temporal signal. Values for *R*^2^ ranged between 0.031 - 0.36 which corresponded again to full genome and gene *NS*_*S*_. The *R*^2^ values for the two datasets of TSWV and 73 orthotospovirus *N* genes were in the range 0.094 - 0.22. BEAST version 1.10.4 (Suchard et al., 2018) on CIPRES Science Gateway (www.phylo.org; Miller et al., 2010) and its associated program BEAUti were used for Bayesian analyses. All the genes, segments and full genome sequences were analyzed under the best-fitting substitution model with site heterogeneity modeled with a Gamma distribution (with four categories) and a fraction of invariable sites, if needed. We used the uncorrelated relaxed clock where each branch in the tree has its own evolutionary rate. Also, we used time-aware GMRF Skyride as a tree prior because we did not have any data on the expansion or population history and it was highly unlikely that constant population size was maintained, chain length of 10^9^, and dates of isolation assigned to each sequence. In the case of the *G*_*N*_*G*_*C*_ gene, we introduced a partition in the alignment, the first half containing the N-terminal 768 amino acids and the second consisting of the C-terminal 821 amino acids. This was done because we found a breakpoint at this position using the program GARD. Each partition had a heterogeneous amino acid substitution model assigned by ProtTest 3. Segments M, S and full genome sequences were also partitioned following the end and the beginning of each respective gene inside the segment. So, segment M was split into three fragments corresponding to genes *NS*_*M*_, *G*_*N*_ and *G*_*C*_, segment S was split into genes *NS*_*S*_ and *N*, while the full genomes were split into genes *N, NS*_*S*_, *NS*_*M*_, *G*_*N*_, *G*_*C*_, and *RdRp*. The respective models selected by ProtTest 3 for each gene were assigned to each partition since the substitution models were unlinked. We used the same parameters as stated above but with unlinked clock models for each partition to account for different rates among branches and with linked trees to obtain a single tree estimated for all the data.

In the phylogeographic analysis of TSWV and 73 orthotospoviruses we focused only on the *N* gene sequences because they greatly outnumbered the rest of the gene sequences available in NCBI. Larger number of sequences gave us higher resolution and more information from different biogeographic realms or ecozones. Assigning ecozones (Scuhultz, 2005) instead of countries to sequences proved to be a better choice since some countries were highly under represented. Furthermore, countries define political entities while ecozones (Fig. 4 and Fig. 5) delimit areas were plants and animals where relatively isolated for long periods of time and therefore share a common evolutionary history. We modeled the sequences with phylogeographic diffusion in discrete space.

All the outputs were analyzed by Tracer 1.7.1 (Rambaut et al., 2018) where convergence and adequate effective sample size (*ESS*) were estimated. To generate the maximum clade credibility (MCC) tree and remove the 10% burn in we used TreeAnnotator 1.10.4. Bayesian posterior probabilities (*BPP*) were used to assess the significance of each node in the MCC tree. FigTree 1.4.3 (tree.bio.ed.ac.uk/software/figtree) was used to visualize all trees and make them publication ready. SpreaD3 was used to prepare the spatiotemporal history and visualize it in a map (Bielejec et al., 2016).

### Assessing the distribution of geographic origins along the MCC phylogenetic tree

Phylogeny-trait association was performed with a software package TreeBreaker (Ansari and Didelot, 2016). The traits assigned were the countries of origin from File S1 for complete genome MCC tree. The analysis was performed on a set of 90,001,000 trees generated with BEAST after removing the 10% burn in.

### *d*_*N*_/*d*_*S*_-based selection analyses

For the analyses of selective constraints operating on the different genes, nucleotide sequences were aligned with MUSCLE (Edgar, 2004) as implemented in MEGA7 (Kumar et al., 2016) using the codon alignment option that preserves open reading frames. Two different methods were used to analyze each codon alignment and inferring differences in selective pressures. The first one was the mixed effects model of evolution (MEME) which uses mixed-effects maximum likelihood approach to test for individual sites that have been subject to episodic positive or diversifying selection (Murrell et al., 2012). The second one was the adaptive branch-site random effects likelihood (aBSREL) model that uses the branch-site models which test for positive selection on a selected proportion of branches (Smith et al., 2015). In both cases the *P* = 0.05 threshold was set for each analysis. These selection analyses were conducted in the www.datamonkey.org server (Weaver et al., 2018).

## SUPPLEMENTARY MATERIAL

Supplementary File S1 (Excel). Orthotospovirus genomic sequence and phylogeographic data.

Supplementary File S2 (Excel). Results of per-site and branch-site selection analyses.

Supplementary Fig. S1 (PDF). Dendrograms of the genomic segments generated with Dendroscope.

Supplementary Fig. S2 (PDF). MCC trees obtained for each genomic segment.

Supplementary Fig. S3 (PDF). Results from the analyses performed with TreeBreaker.

Supplementary Fig. S4 (PDF). Graphical representation of per-site selection analyses. On the abscissa is the nucleotide site and on ordinates the is the likelihood ratio test (LRT) statistic for episodic diversification. Significant sites are those with an LRT value ≥ 4.

## AUTHOR CONTRIBUTIONS

AB and RG collected the data, performed all the analyses and drafted a manuscript. SFE conceived the study, supervised its execution and wrote the manuscript.

## ACKNOWLEDGEMENTS

This work was supported by grant Spain Agencia Estatal de Investigación – FEDER grant BFU2015-65037-P to SFE. AB and RG were supported by Generalitat Valenciana grant GRISOLIAP/2018/005 and by Spain Agencia Estatal de Investigación – FEDER contract BES-2016-077078, respectively.

## REFERENCES

Adams, M. J., Lefkowitz, E. J., King, A. M. Q., Harrach, B. Harrison, R. L., Knowles, N. J., Kropinski, A. M., Krupovic, M., Kuhn, J. H., Mushegian, A. R., Nibert, M., Sabanadzovic, S., Sanfaçon, H., Siddell, S. G., Simmonds, P., Varsani, A., Zerbini, F. M., Gorbalenya, A. E., Davison, A. J., 2017. Changes to taxonomy and the International Code of Virus Classification and Nomenclature ratified by the International Committee on Taxonomy of Viruses (2017). Arch. Virol. 162, 2505–2538.

Ansari, A.M., Didelot, X., 2016. Bayesian inference of the evolution of a phenotype distribution on a phylogenetic tree. Genetics 204, 89–98.

Bielejec, F., Baele, G., Vrancken, B., Suchard, M. A., Rambaut, A., Lemey, P., 2016. SpreaD3: interactive visualisation of spatiotemporal history and trait evolutionary processes. Mol. Biol. Evol. 33, 2167–2169.

Brister, J.R., Ako-Adjei, D., Bao, Y., Blinkova, O., 2015. NCBI viral genomes resource. Nucl. Acids Res. 43, D571–D577.

Darriba, D., Taboada, G.L., Doallo, R., Posada, D., 2011. ProtTest 3: fast selection of best–fit models of protein evolution. Bioinformatics 27, 1164–1165.

Edgar, R.C., 2004. MUSCLE: multiple sequence alignment with high accuracy and high throughput. Nucleic Acids Res. 32, 1792–97.

Eifan, S., Elliot R.M., 2009. Mutational analysis of the Bunyamwera orthobunyavirus nucleocapsid protein gene. J. Virol. 83, 11307–11317.

Fletcher, S. J., Shrestha, A., Peters, J. R., Carroll, B. J., Srinivasan, R., Pappu, H. R., Mitter, N., 2016. The *Tomato spotted wilt virus* genome is processed differentially in its plant host *Arachis hypogaea* and its thrips vector *Frankliniella fusca*. Front. Plant Sci. 7, 1349.

Fu, L., Niu, B., Zhu, Z., Wu, S., Li, W., 2012. CD-HIT: accelerated for clustering the next-generation sequencing data. Bioinformatics 28, 3150–3152.

Gilbertson, R.L., Batuman, O., Webster, C.G., Adkins, S., 2015. Role of the insect supervectors *Bemisia tabaci* and *Frankliniella occidentalis* in the emergence and global spread of plant viruses. Annu. Rev. Virol. 2, 67–93.

Hedil, M., Kormelink, R., 2016. Viral RNA silencing suppression: the enigma of Bunyavirus NS_S_ proteins. Viruses 8, 208.

Hull, R., 2009. Comparative Plant Virology, second edition, Elsevier Academic Press, San Diego CA, USA.

Huson, D.H., Bryant, D., 2006. Application of phylogenetic networks in evolutionary studies. Mol. Biol. Evol. 23, 254–267.

Huson, D.H., Scornavacca, C., 2012. Dendroscope 3: An interactive tool for rooted phylogenetic trees and networks. Syst. Biol. 61, 1061–1067.

Kazutaka, K., Standley, D.M., 2013. MAFFT multiple sequence alignment software version 7: improvements in performance and usability. Mol. Biol. Evol. 30, 772–780.

Knipe, D.M., Howley, P.M., 2013. Fields Virology, sixth edition. Wolters Kluwer/Lippincott Williams & Wilkins Health, Philadelphia PA, USA

Korber, B., 2000. HIV signature and sequence variation analysis, in: Rodrigo, G.A., Learn, G.H. (Eds.), Computational Analysis of HIV Molecular Sequences. Kluwer Academic Publishers, Dordrecht, Netherlands, pp 55–72.

Kosakovsky Pond, S.L., Posada, D., Gravenor, M.B., Woelk, C.H., Frost, S.D., 2006 Automated phylogenetic detection of recombination using a genetic algorithm. Mol. Biol. Evol. 23, 1891–1901.

Kumar S., Stecher G., Tamura K., 2016. MEGA7: molecular evolutionary genetics analysis version 7.0 for bigger datasets. Mol. Biol. Evol. 33, 1870–1874.

Miller, M.A., Pfeiffer, W., Schwartz, T., 2010. Creating the CIPRES science gateway for inference of large phylogenetic trees. Proceedings of the Gateway Computing Environments Workshop (GCE), IEEE, New Orleans LA, USA, pp. 1–8.

Murrell, B., Wertheim, J.O., Moola, S., Weighill, T., Scheffler, K., Kosakovsky Pond, S.L., 2012. Detecting individual sites subject to episodic diversifying selection. PLoS Genet. 8, e1002764.

Nguyen, L.-T., Schmidt, H.A., von Haeseler, A., Minh, B.Q., 2015. IQ-TREE: A fast and effective stochastic algorithm for estimating maximum-likelihood phylogenies. Mol. Biol. Evol. 32, 268–274.

Oliver, J. E., Whitfield, A. E., 2016. The genus *Tospovirus*: emerging bunyaviruses that threaten food security. Annu. Rev. Virol. 3, 101–124.

Pappu, H.R., Jones, R.A., Jain, R.K., 2009. Global status of tospovirus epidemics in diverse cropping systems: successes achieved and challenges ahead. Virus Res. 141, 219–236.

Rambaut, A., Lam, T.T., Max Carvalho, L., Pybus, O.G., 2016. Exploring the temporal structure of heterochronous sequences using TempEst (formerly Path-O-Gen). Virus Evol 2, vew007.

Rambaut, A., Drummond, A.J., Xie, D., Baele, G., Suchard, M.A., 2018. Posterior summarisation in Bayesian phylogenetics using Tracer 1.7. Syst. Biol. 67, 901–904.

Ribeiro, D., Borst, J. W., Goldbach, R., Kormelink, R., 2009. *Tomato spotted wilt virus* nucleocapsid protein interacts with both viral glycoproteins G_N_ and G_C_ in planta. Virology 383, 121–130.

Rotenberg, D., Jacobson, A.L., Schneweis, D.J., Whitfield, A.E., 2015. Thrips transmission of tospoviruses. Curr. Opin. Virol. 15, 80–89.

Scuhultz, J., 2005. The Ecozones of the World, 2^nd^ ed. Springer, New York, NY.

Shalileh, S., Ogada, P.A., Moualeu, D.P., Poehling, H.M., 2016. Manipulation of *Frankliniella occidentalis* (Thysanoptera: Thripidae) by *Tomato spotted wilt virus* (tospovirus) via the host plant nutrients to enhance its transmission and spread. Environ. Entomol. 45, 1235–1242.

Singh, P., Savithri, H., 2015. GBNV encoded movement protein (NS_M_) remodels ER network via C-terminal coiled coil domain. Virology 482, 133–146.

Smith, M.D., Wertheim, J.O., Weaver, S., Murrell, B., Scheffler, K., Kosakovsky Pond, S.L., 2015. Less is more: an adaptive branch-site random effects model for efficient detection of episodic diversifying selection. Mol. Biol. Evol. 32, 1342–1353.

Storms, M.M., Kormelink, R., Peters, D., Van Lent, J.W., Goldbach, R.W., 1995. The nonstructural NS_M_ protein of *Tomato spotted wilt virus* induces tubular structures in plant and insect cells. Virology 214, 485–493.

Suchard, M.A., Lemey, P., Baele, G., Ayres, D.L., Drummond, A.J., Rambaut, A. 2018. Bayesian phylogenetic and phylodynamic data integration using BEAST 1.10. Virus Evol. 4, vey016.

Sundaraj, S., Srinivasan, R., Culbreath, A.K., Riley, D.G., Pappu, H.R., 2014. Host plant resistance against *Tomato spotted wilt virus* in peanut (*Arachis hypogaea*) and its impact on susceptibility to the virus, virus population genetics, and vector feeding behavior and survival. Phytopathology 104, 202–210.

Tsompana, M., Abad, J., Purugganan, M., Moyer, J.W., 2004. The molecular population genetics of the Tomato spotted wilt virus (TSWV) genome: Evolutionary genetics of TSWV. Mol. Ecol. 14, 53–66.

Villesen, P., 2007. FaBox: an online toolbox for FASTA sequences. Mol. Ecol. Notes 7, 965–968.

Walter, C.T., Costa Bento, D.F., Guerrero Alonso, A., Barr, J.N., 2011. Amino acid changes within the *Bunyamwera virus* nucleocapsid protein differentially affect the mRNA transcription and RNA replication activities of assembled ribonucleoprotein templates. J. Gen. Virol. 92, 82–84.

Weaver, S., Shank, S.D., Spielman, S.J., Li, M., Muse, S.V., Kosakovsky Pond, S.L., 2018. Datamonkey 2.0: a modern web application for characterizing selective and other evolutionary processes. Mol. Biol. Evol. 35, 773–777.

